# Cascaded Dual-Targeting (WTW/TAT) Liposomes Delivery DOX for Precision Glioblastoma Treatment

**DOI:** 10.64898/2026.11.02.760854

**Authors:** Xiaoxu Li, Xingxing Gao, Juan Wang, Xiaoming Liao, Zhongbing Huang, Guangfu Yin

## Abstract

Effective chemotherapy for glioblastoma (GBM) is severely restricted by the blood–brain barrier (BBB) and blood–tumor barrier (BTB). To overcome these sequential delivery constraints, we developed a dual-targeted liposomal nanoplatform (WT-LIP) co-modified with a WTW peptide for receptor-mediated BBB transcytosis and GBM targeting, and a cell-penetrating TAT peptide for deep tissue penetration. Doxorubicin-loaded liposomes (DOX@WT-LIP) exhibited significantly enhanced transendothelial transport and downstream tumor cell compared to Angiopep-2/TAT dual-modified liposomes in a BBB/GBM co-culture model. In orthotopic GBM xenografts, WT-LIP demonstrated preferential tumor accumulation (2.16-fold accumulation of tumor-to-normal brain). In vivo, DOX@WT-LIP significantly suppressed tumor progression (74.92% inhibition rate), prolonged survival, and exhibited favorable systemic biosafety. This ligand-guided, cascade-targeting strategy offers a robust approach for penetrating intracranial barriers and improving GBM targeted therapy.

## 1. Introduction

Glioblastoma (GBM) is a highly aggressive brain tumor with a poor prognosis, as current treatments such as surgery and chemotherapy are limited by incomplete tumor resection and frequent recurrence.^1, 2^ Consequently, patients have a median survival of only 8 months and a 5-year survival rate of 7.2%.^3^ This poor prognosis is largely attributed to the blood-brain barrier (BBB), where tight junctions (TJs) between brain capillary endothelial cells (BCECs) restrict the transport of therapeutic agents and other macromolecules.^4–6^ Although GBM disrupts the BBB to form a heterogeneous blood-brain tumor barrier (BBTB) with increased leakage, the BBTB retains key BBB features, including TJs and efflux transporters in BCECs.^7–9^ As a result, drug penetration remains limited, making BBB/BBTB permeability a central challenge in GBM therapy. ^10, 11^

Liposomes are clinically validated nanocarriers that can prolong circulation time and reduce systemic toxicity, exemplified by the first FDA-approved liposomal formulation, Doxil.^12, 13^ Owing to their biodegradability and biocompatibility, appropriately engineered liposomes are promising vehicles for targeted GBM therapy.^14^ Receptor-mediated transport is an effective strategy to promote BBB crossing and improve GBM targeting by engaging overexpressed receptors.^7, 15, 16^ For instance, single-ligand modified D8-sLip/DOX have been developed to target nAChRs on the BBB to improve brain delivery ^17^ or folate-modified liposomal DOX enhanced uptake by folate receptors on GBM cells to enhance tumor uptake.^18^ While dual-ligand liposomes that target receptors on both the BBB and GBM (e.g., OX26/CTX-PL/pC27) have also been explored to further improve delivery,^19–22^ achieving uniform and reproducible ligand density remains technically challenging. Accordingly, increasing attention has shifted to single ligands that can bind receptors expressed on both the BBB and GBM, such as transferrin (Tf) for transferrin receptor (TfR) and angiopep-2 for low-density lipoprotein receptor-related protein (LRP).^23, 24^ However, apparent binding performance of these ligands when receptor binding is efficient, excessively high affinity can hinder ligand-receptor dissociation, promoting lysosomal sequestration and reducing transcytosis across the BBB, thereby limiting drug accumulation in GBM.^25, 26^ Similarly, aniopep-2 can enhance brain penetration, but its broad LRP recognition may increase off-target uptake and yield suboptimal GBM accumulation. These limitations underscore the need for advanced liposome designs that enable efficient BBB transport while maintaining precise GBM targeting.^27–29^

Generally, ligand-functionalized liposomes for GBM delivery undergo a cascaded of steps: (i) specific recognition of receptors on BCEC; (ii) internalization from luminal (the blood-side) membrane into the cytoplasm; (iii) transcytosis and exocytosis at the abluminal (brain-side) membrane, accompanied by dissociation from the BCECs surface; (iv) recognition of receptors on glioblastoma cells (GCs); and (v) internalization into GCs and followed by intracellular drug release.^30, 31^ Therefore, three factors are critical for effective GBM-targeted delivery: ligand specificity toward both BCECs and GCs, membrane/transcytosis competence of the liposomal system, and adaptive ligand–receptor affinity across these two cellular barriers. An ideal dual-targeting ligand should provide selective binding to both BCECs and GCs while exhibiting tunable (context-dependent) binding strength to support BBB transcytosis and subsequent tumor-cell uptake.

In our previous work, we identified a dual-targeting 7-mer peptide (WTW) for GBM using phage display biopanning and affinity–adaptability analysis. The WTW peptide affinity binds to BCECs, enabling transport across the BBB. Upon reaching the brain side, it dissociates and specifically binds to GBM cells, enhancing BBB penetration and tumor accumulation.^32^ The trans-activating (TAT) peptide, a small amphipathic cell-penetrating peptide, can efficiently traverse cell membranes but lacks specificity, leading to off-target effects.^33, 34^ Although TAT-liposomes show a 2.5-fold increase in brain accumulation,^35^ nonspecific distribution remains a concern. Herein, we synthesized the GBM-targeting drug delivery system DOX@WT-LIP (Scheme 1) by considering the cascade process and utilizing functionalized ligands, payloads, and drug release mechanisms for a synergistic effect.^36, 37^ Our DOX@WT-LIP achieves highly efficient BBB transcytosis and enhanced GBM accumulation through the moderate affinity of its WTW ligand for BCECs and strong affinity for GBM cells, preventing lysosomal capture and promoting targeted drug release. Enhanced by the synergistic effect of WTW and TAT, alongside liposome properties ensuring prolonged circulation, this regulated cascade delivery significantly improves GBM therapeutic efficacy.

**Schematic 1.**
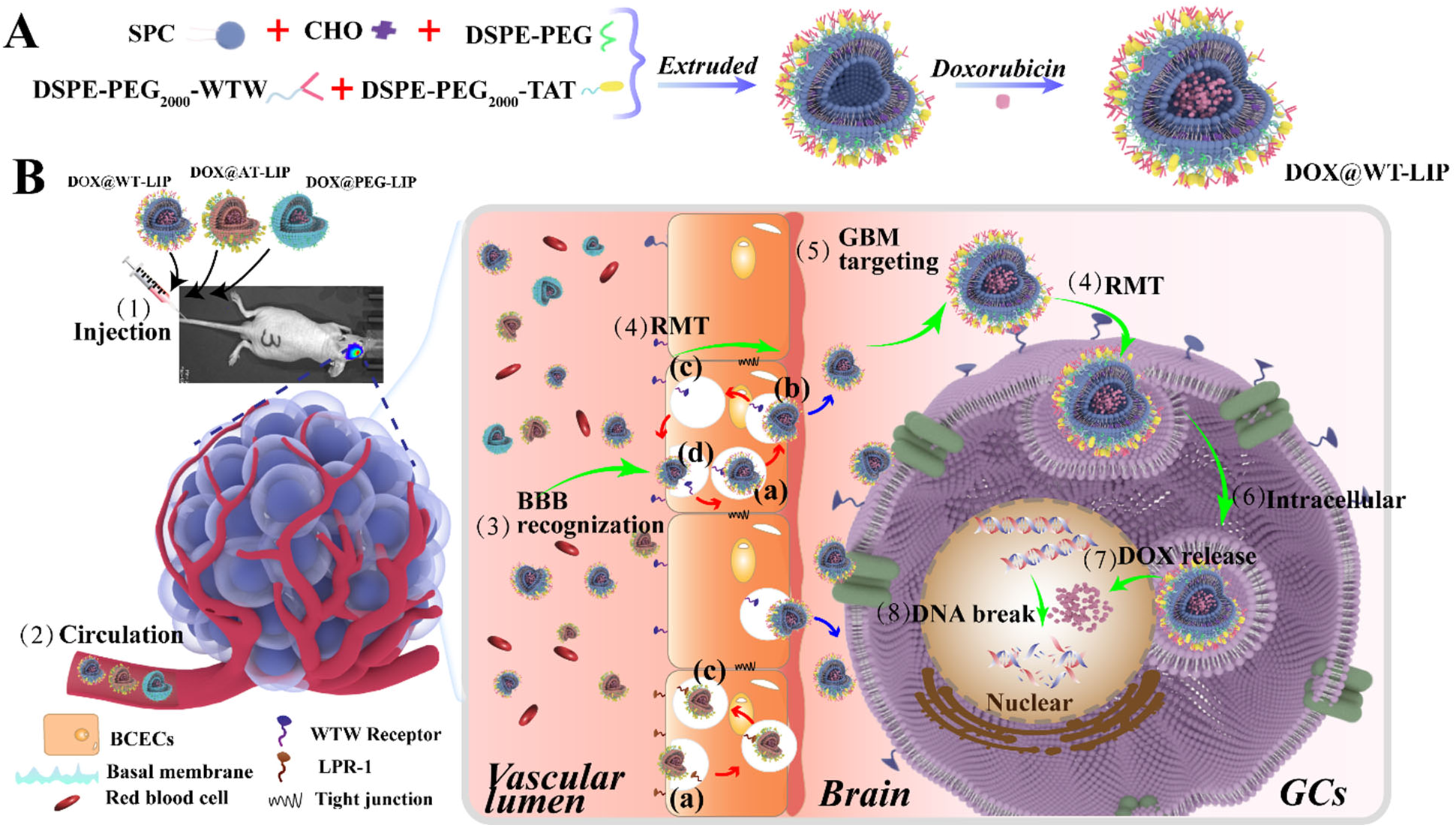
Schematic of synthesis of DOX@WT-LIP and its mechanism for aggregation to GBM through BBB transport. (A). Synthesis method of DOX@WT-LIP; (B). Illustration of DOX@WT-LIP programmed to recognize receptors on BBB and then target receptors on GBM cells after transcytosis, then releasing DOX for therapeutic effect. After (1) intravenous injection and (2) circulation in the bloodstream, (3) DOX@WT-LIP recognize and bind to on receptors on endothelial of BBB, and (4) transports into the brain through receptors circulation, including (a) internalization by BCECs receptors, (b) dissociation from BCECs receptors and release into brain parenchyma, (c) receptors recycling back to BCECs surface, (d) subsequent DOX@WT-LIP bind to the receptors (red arrows represented receptors recycling). Following BBB traversal, (5) DOX@WT-LIP targets GBM receptors and (6) is subsequently internalized via RMT. Once inside the GBM cells, DOX is released, leading to DNA double-strands breaks and subsequent apoptosis of GBM cells (green arrows represented the cascade process delivery process).

## 2. Resutles

### 2.1 Preparation and characteristics of peptides-modified liposomes

The WTW peptide was conjugated to PEG_2000_ via a Michael addition reaction between its sulfhydryl group and the terminal maleimide of DSPE-PEG_2000_-Mal (Figure 1A). DSPE-PEG_2000_-Ang and DSPE-PEG_2000_-TAT were synthesized similarly. The successful conjugation and structure of these functionalized DSPE-PEG_2000_ constructs were confirmed by proton Nuclear Magnetic Resonance (^1^H-NMR, Figure 1B, Figure S1). Doxorubicin-loaded liposomes modified with both WTW and TAT (DOX@WT-LIP) were then synthesized by combining SPC, CHOL, DSPE-PEG_2000_, DSPE-PEG_2000_-TAT, and DSPE-PEG_2000_-WTW (Table S1). Doxorubicin (DOX) was actively encapsulated using the classical transmembrane pH gradient method,^38–40^ achieving a high DOX loading efficiency (LE%) of 2.84 ± 0.08% and an encapsulation efficiency (EE%) of 94.17 ± 1.30% in DOX@WT-LIP, with similar efficiencies observed in its derivatives (Table S2). Dynamic Light Scattering (DLS) analysis showed hydrodynamic diameters of 118.93 ± 0.41 nm for DOX@AT-LIP, 111.27 ± 0.85 nm for DOX@WT-LIP, and 98.86 ± 0.66 nm for DOX@PEG-LIP, with corresponding ζ-potentials of -20.57 ± 0.78 mV, -21.07 ± 0.62 mV, and -3.58 ± 0.48 mV, respectively (Figure 1C-F). The larger size of DOX@AT-LIP is probably attributed to the higher molecular weight of Angiopep-2 (2.35 kDa) compared to WTW (1.12 kDa). Transmission Electron Microscopy (TEM) confirmed well-defined nanospheres for DOX@WT-LIP (38.25 ± 4.97 nm) and its derivatives (DOX@AT-LIP: 44.31 ± 5.96 nm, DOX@PEG-LIP was 27.70 ± 4.00 nm), while cryo-Electron Microscopy (cryo-EM) additionally revealed their internal single-chambered vesicular structure (Figure 1G/H, Figure S1B).

**Figure 1.**
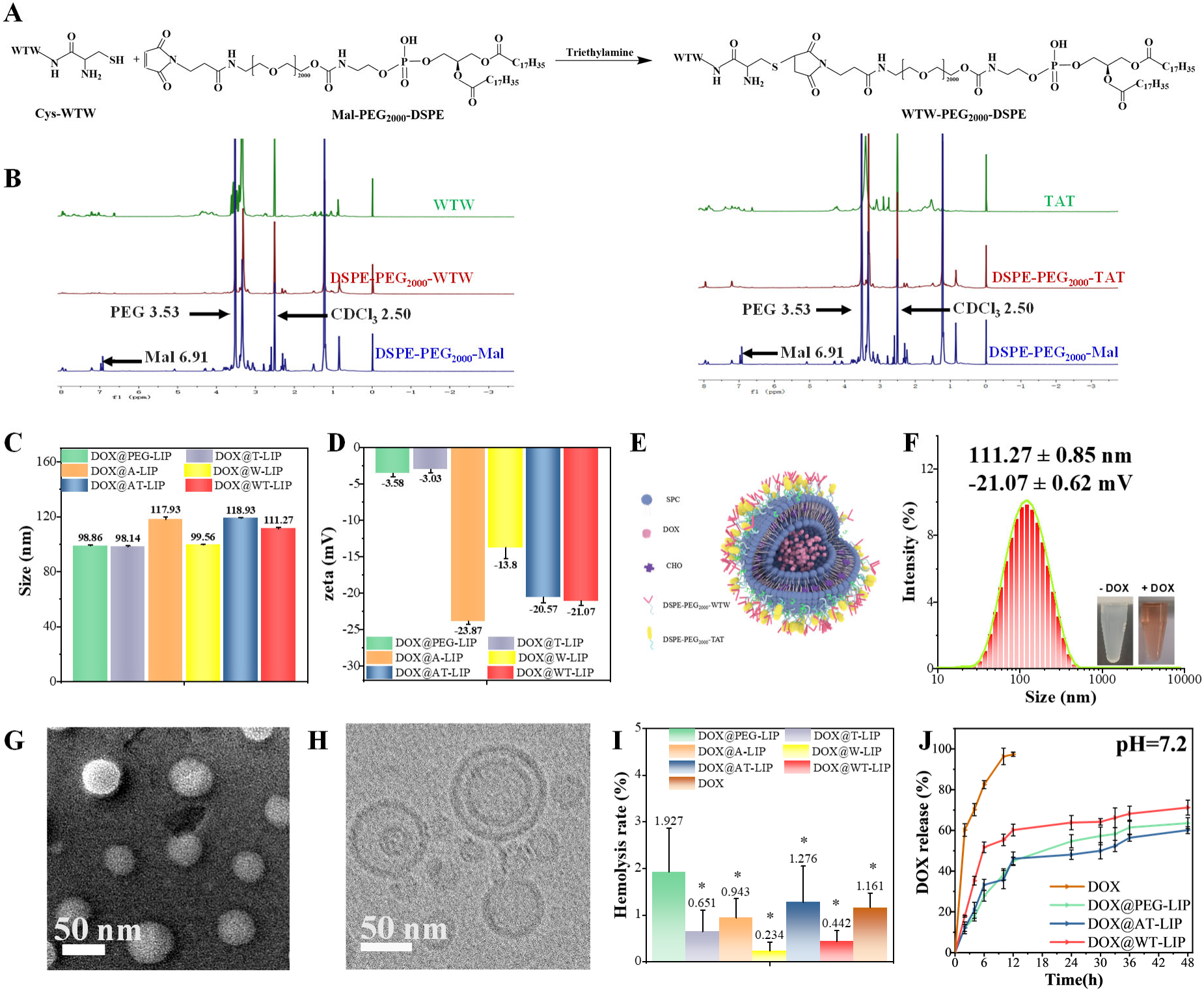
Preparation and characteristics of peptides-modified liposomes. (B) ^1^H-NMR spectra of WTW, TAT, DSPE-PEG_2000_-Mal and DSPE-PEG_2000_-WTW/TAT; (C) Hydrodynamic diameters size and (D) zeta-potential of DOX@WT-LIP and it derives; (E)Scheme structure of DOX@WT-LIP and (F) its size distribution by DLS. (G) Transmission electron microscopy (TEM) image and (H) Cryogenic electric microscopy (cryo-EM) image of DOX@WT-LIP; (I) Hemocompatibility test of DOX-loaded liposomes (J) *In vitro* drug release from different DOX-loaded liposomes in 48 h; \**p* < 0.05

### 2.2 DOX release behavior and Hemocompatibility

Glioblastoma target delivery system requires prolonged stable circulation to increase the probability of binding to the target cells, drug releasing behavior and hemocompatibility is the prerequisite of their safe application in circulatory systems.^41, 42^ In vitro drug release from DOX-loaded liposomes was assessed under sink conditions using a dialysis method. As shown in Figure 1I, free DOX is fully diffused within 10 hours. In contrast, all liposomal formulations exhibited gradual, sustained release over 48 hours, with cumulative release of 60-71%. The lack of an initial burst release suggests that DOX was effectively encapsulated within the liposomal vesicles and colloidal salt deposits via the pH gradient method.^43–45^ Hemolysis assays showed that all liposomal formulations had hemolysis ratio were less than 2% and obviously below the 5% safety threshold. Compared with other groups, the hemolysis ratios of DOX@WT-LIP (0.44 ± 0.23%) were the lowest, DOX@AT-LIP (1.28 ± 0.78% for DOX@AT-LIP) was followed, and the highest of that was DOX@PEG-LIP (1.93 ± 0.94%) which indicated that DOX@WT-LIP possessed a good hemocompatibility (Figure 1J).

### 2.3 Cellular uptake

U251-MG (mice GBM cell), bEnd.3 (BBB-related cell), M1800-57 (mice astrocytes cell), HUVEC (Human umbilical vein endothelial cell), L929 (mouse fibroblast cell), and RAW 264.7 (mouse macrophage cell) cells were incubated with Coumarin-6 (Co6)-labeled WT-LIP, AT-LIP, and PEG-LIP for 24 hours and subsequently observed using confocal laser scanning microscopy (CLSM). The CLSM and quantification results (Figure 2A/B) demonstrated that the fluorescence intensity of Co6-WT-LIP in U251-MG and bEnd.3 was significantly higher than in cells incubated with Co6-AT-LIP and Co6-PEG-LIP. Notably, the uptake of Co6-WT-LIP by U251-MG and bEnd.3 cells was also significantly higher compared to M1800-57, HUVEC, L929, and RAW 264.7 cells. While the fluorescence intensity of Co6-AT-LIP in bEnd.3 was higher than that of Co6-PEG-LIP, it was significantly lower than Co6-WT-LIP. To further quantify the relative affinities, the tumor-to-BBB uptake ratio (T_U/B_) was calculated as the fluorescence intensity ratio of Co6-labeled liposomes in U251-MG to bEnd.3 cells. As shown in Figure 2C, the T_U/B_ of Co6-WT-LIP was 2.04 ± 1.27, which was notably higher than that of Co6-AT-LIP (1.74 ± 0.81). These results collectively indicate that Co6-WT-LIP exhibits superior selective binding for cellular uptake by U251-MG and bEnd.3 cells, highlighting its significant potential to target BCECs and GCs. Although Co6-AT-LIP also showed selective cellular uptake compared with Co6-PEG-LIP, its selectivity was considerably lower than that of Co6-WT-LIP. The large T_U/B_ value of Co6-WT-LIP is particularly advantageous, as it is conducive to its dissociation from the BBB and subsequent aggregation in GBM after penetrating the brain, from the perspective of affinity compatibility.

**Figure 2.**
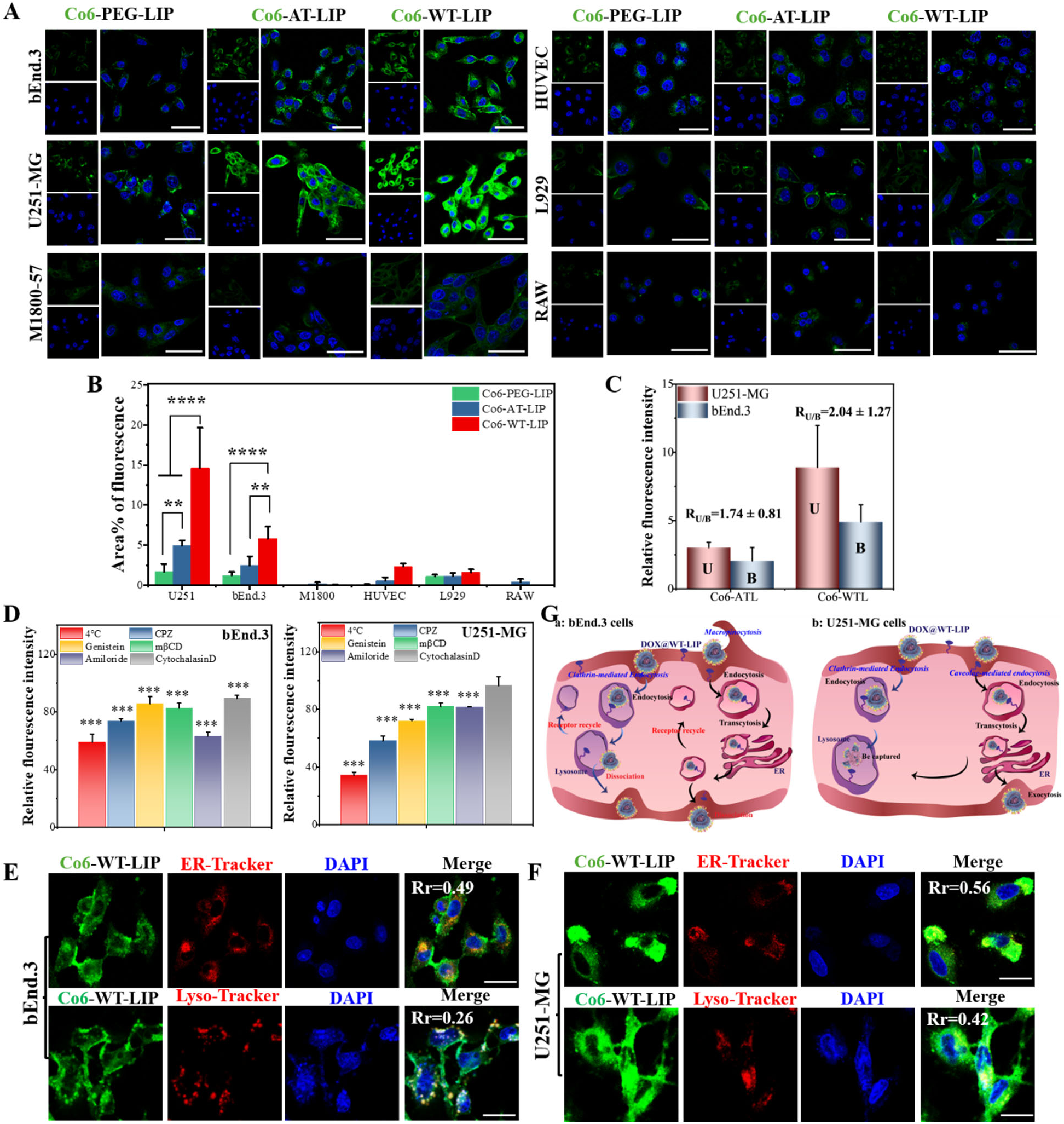
Cellular uptake, endocytic behavior, and lysosome/ER colocalization. (A). The fluorescence image of Co6-PL, Co6-ATL, and Co6-WTL on different cells; (B). The quantification of area% fluorescence of Co6-loaded liposomes in different cells from A; (C). Relative affinity (R_U/B_) of liposomes to U251-MG and bEnd.3 cells; (D) The inhibiting effects of endocytic inhibitors and low temperature on cellular uptake of U251-MG and bEnd.3 cells; The lysosome/ER colocalization of bEnd.3 (E) and U251-MG (F) cells, bars represent 100 µm; (G) Scheme diagram of DOX@WT-LIP endocytic behavior in bEnd.3 and U251-MG cells; ** p < 0.01, *** p < 0.001, and **** p < 0.0001

### 2.4 Endocytic behavior of Co6-WT-LIP

To identify the endocytic pathways responsible for Co6-WT-LIP uptake, bEnd.3 and U251-MG cells were pretreated with chlorpromazine (CPZ; clathrin-mediated endocytosis inhibitor), genistein (caveolae-mediated endocytosis inhibitor), amiloride (macropinocytosis inhibitor; associated with Na^+^/H^+^ exchange and PI3K-dependent processes), methyl-*β* -cyclodextrin (m*β*CD; cholesterol depletion/inhibition of lipid raft–associated uptake), cytochalasin D (actin polymerization inhibitor; inhibits macropinocytosis), or incubated at 4 °C to suppress energy-dependent internalization.

In U251-MG cells (Figure 2D), Co6-WT-LIP uptake decreased to 41.9% with CPZ, 28.0% with genistein, and 65.9% at 4 °C, whereas the other inhibitors produced only minor effects. These results suggest that uptake of U251-MG cells is predominantly energy-dependent and proceeds mainly through clathrin-mediated and caveolae-mediated endocytosis. In bEnd.3 cells (Figure 2D), Co6-WT-LIP uptake decreased to 36.9% with amiloride, 26.5% with CPZ, and 41.1% at 4 °C, while the other inhibitors had little effect, indicating that internalization in bEnd.3 cells is primarily mediated by macropinocytosis (likely promoted by the TAT peptide) together with clathrin-mediated endocytosis.

Clathrin- and caveolae-mediated pathways are common mechanisms of receptor-mediated endocytosis, whereas TAT is known to promote nonselective macropinocytosis.^46^ Nanomedicines internalized via clathrin-mediated endocytosis are often trafficked to lysosomes, whereas caveolae-mediated uptake can favor trafficking routes that avoid lysosomal degradation and facilitate transcellular transport.^47–49^ In contrast, cargo internalized by macropinocytosis may either recycle back to the plasma membrane or traffic to lysosomes depending on vesicle maturation.^50, 51^ Therefore, the intracellular fate of Co6-WT-LIP after uptake by bEnd.3 and U251-MG cells warrants further investigation through colocalization and trafficking studies.

### 2.5 Lysosome/ER colocalization of Co6-WTL

The intracellular trafficking of Co6-WT-LIP in U251-MG and bEnd.3 cells was examined by confocal laser scanning microscopy (CLSM). The extent of colocalization between Co6-WT-LIP and the endoplasmic reticulum (ER) or lysosomes was quantified using Pearson’s correlation coefficient (*R*_r_) in ImageJ. The ER and lysosomes were stained with ER-Tracker and Lyso-Tracker, respectively. In bEnd.3 cells (Figure 2E), Co6-WT-LIP showed stronger colocalization with the ER (*R*_r_ = 0.64 ± 0.49) but weak colocalization with lysosomes (*R*_r_ = 0.26 ± 0.06), indicating that Co6-WT-LIP is predominantly distributed in the ER/cytoplasm and is less extensively sorted to lysosomes. This behavior aligns with the endocytosis results in bEnd.3 cells, where macropinocytosis (likely promoted by TAT) and clathrin-mediated endocytosis contributed to uptake. In U251-MG cells (Figure 2F), Co6-WT-LIP exhibited appreciable colocalization with both ER-Tracker and Lyso-Tracker signals, with *R*_r_ = 0.56 ± 58 for ER and *R*_r_ = 0.42 ± 46 for lysosomes. These results suggest that a substantial fraction of internalized Co6-WT-LIP enters the endo/lysosomal pathway, while another fraction traffics to the ER. This distribution is consistent with the inhibitor studies, which indicated that uptake in U251-MG cells proceeds mainly through clathrin-mediated and caveolae-mediated endocytosis.

Peptide-modified liposomes can undergo receptor-mediated internalization, intracellular sorting, and subsequent exocytosis.^52, 53^ Importantly, ligand–receptor affinity can influence post-entry trafficking and transcytosis efficiency.^9, 54^ High-affinity interactions may favor clathrin-mediated uptake and increase lysosomal sequestration, thereby limiting transcytosis.^52, 55, 56^ In addition, TAT can interact with negatively charged membrane components and promote cellular entry, which may facilitate intracellular trafficking routes that support membrane penetration.^57–60^ Based on the colocalization results (Figure 2A/B), we speculate that in U251-MG cells a portion of Co6-WT-LIP internalized via clathrin-mediated endocytosis is directed to lysosomes, whereas another portion entering via caveolae-mediated uptake can traffic toward the ER and potentially support onward transport (Figure 2G). In bEnd.3 cells, the limited lysosomal colocalization suggests that Co6-WT-LIP may partially avoid lysosomal routing after internalization, which could be favorable for exocytosis/transcytosis (Figure 2G). However, cellular uptake and intracellular localization alone do not demonstrate BBB crossing efficiency. Therefore, we next evaluated transcytosis using an in vitro BBB model.

Liposomes internalized via clathrin-mediated endocytosis are often trafficked to lysosomes,^48^ and macropinosomes formed during macropinocytosis can also mature and fuse with late endosomes/lysosomes.^53, 61^ In bEnd.3 cells (Figure 2D-G), Co6-WT-LIP entered through clathrin-mediated endocytosis and macropinocytosis, yet showed limited colocalization with Lyso-Tracker, suggesting partial escape from lysosomal routing and subsequent exocytosis. This behavior may arise from WTW dissociation from its receptor, enabling repeated binding/uptake at the endothelial surface. Angiopep-2 binds LRP-1 with high affinity on both glioma cells and BCECs.^62, 63^ Such strong binding can hinder dissociation, promote lysosomal sequestration, and reduce receptor recycling,^54^ potentially explaining the lower uptake of Co6-AT-LIP in bEnd.3 cells. However, uptake and trafficking do not directly predict BBB crossing; therefore, we next evaluated transcytosis in an in vitro BBB model.

### 2.6 Cytotoxicity, ***γ***-H2AX expression, and cell ultrastructure

For cytotoxicity, U251-MG cells were incubated with free DOX, DOX@PEG-LIP, DOX@AT-LIP, or DOX@WT-LIP for the indicated times, and viability was quantified by MTT assay (Figure 3A). In U251-MG cells, all formulations showed dose-dependent cytotoxicity at 24 and 48 h. At 48 h, DOX@WT-LIP L produced significantly stronger inhibition than DOX@PEG-LIP and DOX@AT-LIP at 5–20 μg/mL. The IC_50_ values at 48 h were 3.75 μg/mL (free DOX), 5 μg/mL (DOX@WT-LIP), 10 μg/mL (DOX@AT-LIP), and 12 μg/mL (DOX@PEG-LIP).

**Figure 3.**
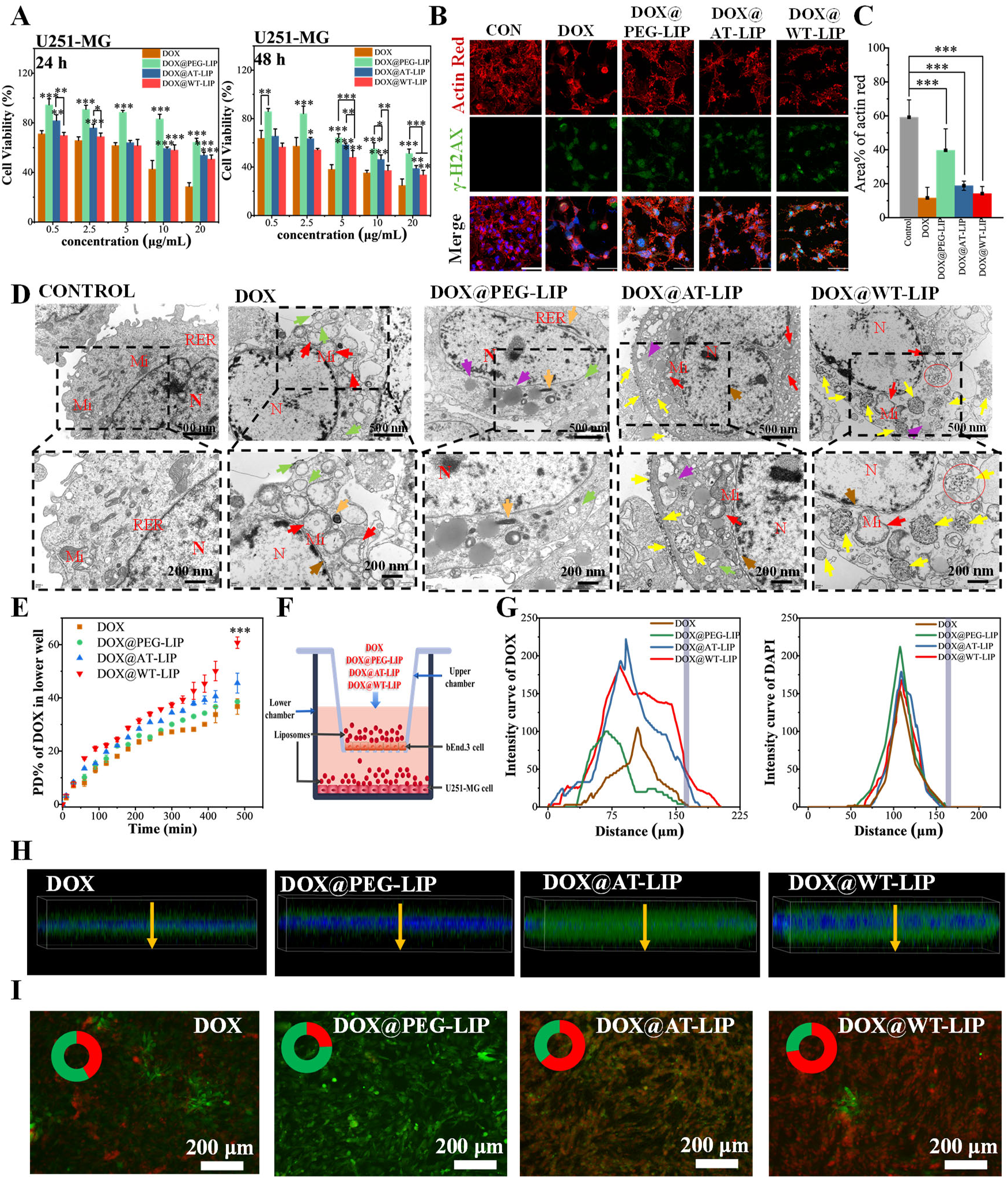
Tumor inhibition, cellular ultrastructure, γ-H2AX expression and evaluation of liposomes *in vitro* BBB transcytosis. (A) The MTT assay of free DOX, DOX@WT-LIP and it derives against U251-MG cells for 24 h, and 48 h; Expression (B) and fluorescence quantitative (C) of γ-H2AX and in U251-MG cells treated with liposomes for 24 h, bars represent 100 µm; (D) TEM image of U251-MG cells from the control group, free DOX, DOX@PEG-LIP, DXO@AT-LIP and DOX@WT-LIP group; (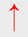: mitochondrial swelling, 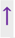: lipid droplets, 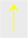: material-like substances, 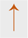: widening of perinuclear gap, 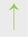: expansion of the RER, 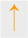: mitochondrial consolidation, ○:endocytosis, Mi: mitochondria, N: nuclear, RER: rough endoplasmic reticulum); (E) PD% of total DOX reaching the lower well after introducing free DOX, DOX@PEG-LIP, DXO@AT-LIP and DOX@WT-LIP group into bEnd.3 monolayers at different time intervals; (F) Schematic of the *in vitro* BBB transcytosis experiments; (H/I) Results of the mean fluorescence intensity of U251-MG cells in lower chambers after free DOX, DOX@PEG-LIP, DXO@AT-LIP and DOX@WT-LIP group incubation in upper chambers for 24 h; (G) Intensity curve of DOX at the indicative location and distance from H and Intensity curve of cytoskeletal actin at the indicative location and distance from H; (H) 3D confocal images of bEnd.3 monolayers in the upper chamber after incubation with free DOX, DOX@PEG-LIP, DXO@AT-LIP and DOX@WT-LIP for 24 h, yellow profile line represents the location and distance for quantitative analysis of fluorescence signal; #*p*< 0.001; (I) Live/Dead fluorescence of U251-MG cells in lower chamber after free DOX, DOX@PEG-LIP, DXO@AT-LIP and DOX@WT-LIP incubation in upper chamber for 24 h. \**p* < 0.05, \*\**p* < 0.01, and \*\*\**p* < 0.001

Doxorubicin (DOX) intercalates into DNA and inhibits topoisomerase II–mediated repair, causing DNA double-strand breaks (DSBs) and phosphorylation of histone H2AX (*γ*-H2AX), a sensitive marker of DNA damage.^64^ DOX can also remodel the actin cytoskeleton, disrupt central actin fibers, and weaken cell adhesion, thereby promoting cell detachment.^65, 66^ After 24 h, free DOX, DOX@AT-LIP, and DOX@WT-LIP induced pronounced morphological changes in U251-MG cells, with cells transitioning from a cuboidal morphology to a shrunken phenotype with spindle-like actin structures and disrupted nuclei (Figure 3B-C). In contrast, the DOX@PEG-LIP group showed comparatively weaker morphological changes and less reduction in cell density.

For cell ultrastructure observation, U251-MG cells were examined by transmission electron microscopy (TEM) after 48 h treatment to assess ultrastructural changes and intracellular behavior of DOX-loaded liposomes. Untreated cells showed intact morphology, including regular nuclei, evenly distributed chromatin, and well-preserved organelles (mitochondria, ER, and Golgi) (Figure 3D). In the free DOX group, severe damage was observed, including nuclear rupture, membrane disruption, cytoplasmic leakage, chromatin dissolution, mitochondrial swelling, and dilation of the rough ER. Similar severe disruption was observed in the DOX@WT-LIP group, where many cells exhibited chromatin dissolution, membrane rupture, cytoplasmic loss, and mitochondrial swelling. Notably, membrane-bound nanoparticle-like structures (∼50–80 nm) were detected in the cytoplasm (red circles), consistent with the size of DOX@WT-LIP measured by TEM and suggesting efficient cellular internalization. In the DOX@AT-LIP group, similar particulate structures were occasionally observed, but clear endocytic vesicles were not evident; overall, cells showed moderate injury and some retained relatively intact morphology. By contrast, the DOX@PEG-LIP group exhibited the mildest ultrastructural disruption, with cellular architecture largely preserved.

Liposomes typically enter cells through receptor-associated uptake processes involving ligand binding, endocytosis, and intracellular drug release.^9, 67^ The abundant intracellular particles in the DOX@WT-LIP group support efficient internalization. The WTW ligand likely promotes binding to GPCRs on U251-MG cells, increasing liposome association with the cell surface, while the TAT peptide may further enhance membrane penetration and uptake. Following internalization, DOX@WT-LIP is expected to traffic through endocytic pathways (including clathrin-mediated routes) and undergo lysosomal processing, facilitating DOX release and amplifying DNA damage and cytotoxicity in GBM cells. In contrast, although angiopep-2 in DOX@AT-LIP can bind LRP-1 and mediate cellular uptake, comparatively weaker accumulation in U251-MG cells may limit intracellular DOX delivery and reduce efficacy.

### 2.7 Transcytosis of BBB/GBM model *in vitro*

In vitro BBB monolayer model was established by seeding bEnd.3 cells in upper chamber of 12-well transwell plate for 14 days and forming a resistant transmembrane (TEER was ∼185 Ω.cm^2)^. To evaluate BBB transcytosis, free DOX or DOX-loaded liposomes were added to the apical (upper) chamber and incubated for 480 min. DOX in the basolateral (lower) chamber was quantified every 30 min. As shown in Figure 3E, the percentage of permeated dose (PD%) increased over time for all groups, and the difference between DOX@WT-LIP and the other formulations gradually widened. At 480 min, DOX@WT-LIP achieved a PD% of 60.74 ± 2.14%, significantly higher than DOX@AT-LIP (45.55 ± 6.74%), DOX@PEG-LIP (38.57 ± 0.64%), and free DOX (36.76 ± 2.92%). These results indicate that DOX@WT-LIP exhibits higher in vitro BBB transcytosis efficiency than its derivates and free DOX due to the mediated effect of dual-targeting peptide WTW.

To further assess post-BBB delivery to tumor cells, an in vitro BBB/GBM co-culture model was constructed by seeding U251-MG cells in the lower chamber after the bEnd.3 monolayer had matured for 14 days (Figure 3F). Free DOX, DOX@PEG-LIP, DOX@AT-LIP, and DOX@WT-LIP were added to the upper chamber and incubated for 24 h. Among all groups, DOX@WT-LIP showed the strongest DOX fluorescence and deepest penetration through the endothelial layer. Quantification of mean fluorescence intensity across the bEnd.3 monolayer (apical to basolateral) showed penetration depths of 160 μm (DOX@PEG-LIP), 178 μm (DOX@AT-LIP), and 202 μm (DOX@WT-LIP). Free DOX (MW 579 Da) also diffused across the monolayer with a measured depth of 163 μm (Figure 3G). Nuclear (blue) fluorescence profiles were similar across groups (all reaching ∼160 μm), indicating comparable cell distribution through the reconstructed stacks (Figure 3G). The bEnd.3 monolayer was imaged by CLSM and reconstructed in z-stacks (Figure 3H). Overall, DOX@WT-LIP demonstrated greater endothelial penetration and higher DOX signal, suggesting more efficient transport across the bEnd.3 barrier.

U251-MG viability after transport was evaluated using a Live/Dead assay (live: green; dead: red). As shown in Figure 3I, DOX@WT-LIP produced the highest proportion of dead cells, followed by DOX@AT-LIP and free DOX, whereas DOX@PEG-LIP showed the weakest effect. This pattern is consistent with enhanced post-BBB accumulation of DOX@WT-LIP in U251-MG cells and increased DOX-associated cytotoxicity. Collectively, together with the cellular uptake data (Figure 2A/B), these results support that WTW promotes BBB transcytosis and improves subsequent tumor-cell delivery. TAT may contribute to macropinocytosis and intracellular trafficking that reduces lysosomal sequestration, thereby facilitating transport from the luminal to the abluminal side. After BBB crossing, DOX@WT-LIP may preferentially accumulate in glioma cells, enhancing internalization and DOX-mediated cytotoxicity. In contrast, DOX@AT-LIP may undergo stronger retention and/or lysosomal routing in endothelial cells following LRP-1 engagement, which could reduce effective delivery to the tumor compartment.

### 2.8 Biodistribution of DiD-labeled liposomes and pharmacokinetics

Biodistribution of DiD-labeled liposomes were evaluated in U251-luc GBM–bearing mice using an IVIS Spectrum imaging system to assess brain delivery and tumor targeting. DiD–PEG-liposome (DiD-PEG-LIP), DiD–Ang–TAT-liposome (DiD-AT-LIP), and DiD–WTW–TAT-liposome (DiD-WT-LIP) were prepared as previously described. As shown in Figure 4A, DiD-PEG-LIP fluorescence was broadly distributed throughout the body from 0 to 24 h after injection, with no obvious enrichment. DiD-AT-LIP showed increased fluorescence in the chest and brain regions at 2 h, consistent with rapid systemic distribution, followed by a gradual decline; a weak but detectable brain signal remained at 24 h. In contrast, DiD-WT-LIP exhibited a progressive increase in brain-associated fluorescence over time and a markedly stronger brain signal at 24 h relative to other regions, except the cardiac area.

**Figure 4.**
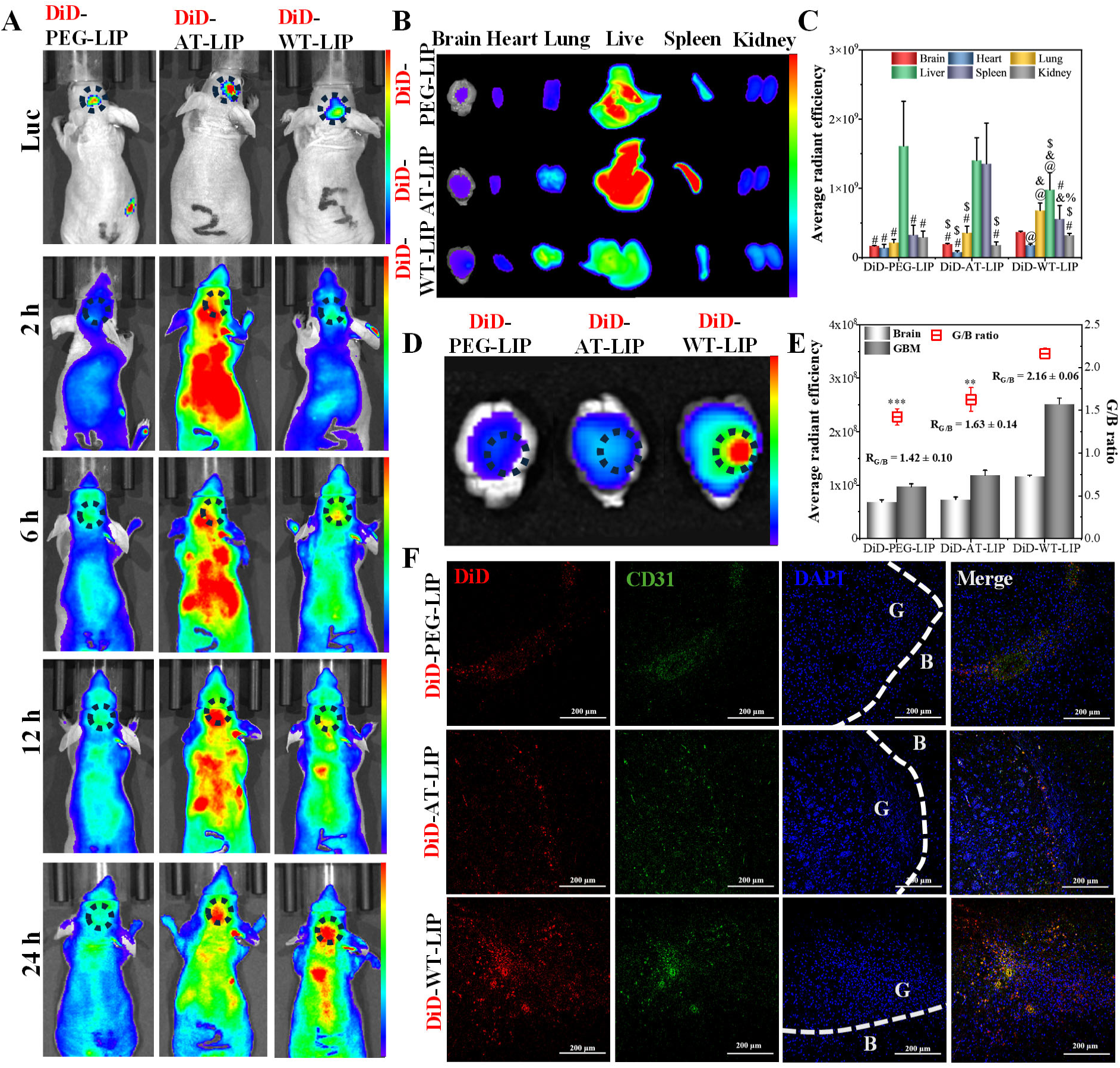
*In vivo* biodistribution of DiD-labeled liposomes. (A) The living image of U251-luc GBM-bearing mice after intravenous injection with DiD-PL, DiD-ATL, and DiD-WTL at different time intervals, the bar represents radiant efficiency from 4.0 × 10^8^ to 2.0 × 10^9^ [p s^−1^ cm^−2^ sr^−1^]/ [µW cm^−2^]; (B) *Ex vivo* imaging of major organs collected at 24 h after injection, bar represents radiant efficiency from 8.0 × 10^7^ to 3.0 × 10^8^ [p s^−1^ cm^−2^ sr^−1^]/ [µW cm^−2^]; (C) Semi-quantitative data of fluorescence signal of organ from D; @: Significant differences with fluorescence signal in brain, *p* <0.05; &: Significant differences with fluorescence signal in heart, *p* <0.05; #: Significant differences with fluorescence signal in lung, *p* <0.05; $: Significant differences with fluorescence signal in lung, *p* <0.05; #: Significant differences with fluorescence signal in liver, *p* <0.05; %: Significant differences with fluorescence signal in spleen, *p* <0.05; (D) *Ex vivo* imaging of brains collected at 24 h after injection, the bar represents radiant efficiency from 1.0 × 10^8^ to 2.0 × 10^9^ [p s^−1^ cm^−2^ sr^−1^]/ [µW cm^−2^]; (E) Left column represented semi-quantitative data of fluorescence signal at the indicative site from B, right column represented G/B ratio of different DiD-labeled liposomes; (F) Fluorescence distribution of DiD-WTL at GBM site after intravenous administration for 24 h, the brain slices were immune-stained with anti-CD31 antibody, bars represent 200 μm. \*\**p* < 0.01, \*\*\**p* < 0.001

Because the liver and spleen are primary organs of the mononuclear phagocyte system (MPS) responsible for nanoparticle clearance, the liver- and spleen-dominant distributions of DiD-PL and DiD-AT-LIP suggest substantial MPS uptake, which may shorten circulation time and reduce brain exposure.^68^ By comparison, DiD-WT-LIP exhibited higher brain accumulation with relatively lower liver and spleen signals, suggesting reduced MPS sequestration and prolonged circulation. This interpretation is consistent with the lower association of Co6-WT-LIP with RAW 264.7 macrophages in uptake studies and the sustained in vivo brain signal of DiD-WT-LIP at 24 h. The elevated lung signal observed for DiD-WT-LIP may be attributable to serum protein adsorption and the resulting increase in apparent particle size (approximately 15 nm in the serum stability test), which could enhance retention in Major organs were collected for ex vivo imaging and semi-quantitative analysis (Figure 4B/C). DiD-PEG-LIP showed predominant accumulation in the liver, whereas DiD-AT-LIP displayed high signals in both liver and spleen. DiD-WT-LIP showed moderate fluorescence in liver, lung, and spleen, with comparatively higher signal in the brain, further formulation optimization may reduce lung-associated accumulation.

Notably, DiD-WT-LIP showed higher fluorescence at the tumor region than DiD-AT-LIP and DiD-PEG-LIP, indicating enhanced accumulation at the GBM site (Figure 4D). Tumor selectivity was quantified by the GBM-to-brain (G/B) ratio, defined as fluorescence at the GBM site divided by fluorescence in normal brain parenchyma. DiD-WT-LIP achieved a G/B ratio of 2.16 ± 0.06, higher than DiD-AT-LIP ( 1.63 ± 0.14 ) and DiD-PEG-LIP ( 1.42 ± 0.10 ) (Figure 4E). Brain sections were further analyzed by immunostaining with anti-CD31, an endothelial marker of brain microvascular endothelial cells. As shown in Figure 4F, DiD-WT-LIP produced stronger fluorescence in brain tissue than DiD-PEG-LIP and DiD-AT-LIP. Moreover, DiD-WT-LIP exhibited higher fluorescence in the GBM region than in normal brain parenchyma, consistent with the elevated G/B ratio, suggesting improved tumor-associated retention/uptake after brain entry. In contrast, DiD-PEG-LIP and DiD-AT-LIP showed weaker tumor-region fluorescence and smaller tumor-to-normal differences, potentially due to lower BBB transport and reduced tumor selectivity.

### 2.9 Efficacy evaluation

For pharmacokinetic evaluation, plasma DOX concentrations were measured in Sprague–Dawley rats after tail-vein injection of DOX@WT-LIP (DOX dose: 4 mg/kg). The plasma profile showed *C*_max_ = 14.148 mg/L, *T*_max_ = 1 h, *t*_1/2_ = 1.965 h, MRT = 14.629 h, and AUC = 52.147 mg·h/L (Table S3 and Figure S3), indicating sustained systemic exposure of DOX delivered by DOX@WT-LIP, which may increase drug availability for accumulation at the GBM site.

An orthotopic U251-luc GBM mouse model was used to compare the efficacy of DOX@PEG-LIP, DOX@AT-LIP, DOX@WT-LIP, and free DOX (Figure 5A). Tumor burden was monitored by IVIS bioluminescence imaging after intraperitoneal injection of D-luciferin potassium salt (150 mg/kg), and signals were semi-quantitatively analyzed. Free DOX and DOX@PEG-LIP induced only modest tumor growth, whereas DOX@WT-LIP produced a more pronounced reduction in luciferase bioluminescence (Figure 5B). Consistently, DOX@WT-LIP achieved a tumor inhibition rate of 74.92%, higher than DOX@AT-LIP (48.58%), DOX@PEG-LIP (37.2%), and free DOX (21.21%) (Figure 5C). In survival analysis, all mice in the DOX@WT-LIP group survived throughout the observation period. In contrast, two of five mice in the free DOX group died during treatment (day 14), and one of five mice in the DOX@AT-LIP group died on day 28, which may be associated with systemic toxicity and/or limited tumor selectivity. Body weight decreased across all treatment groups over 28 days; however, weight loss was less pronounced in the DOX@WT-LIP group (Figure 5D), suggesting a comparatively lower treatment-associated burden under these conditions.

**Figure 5.**
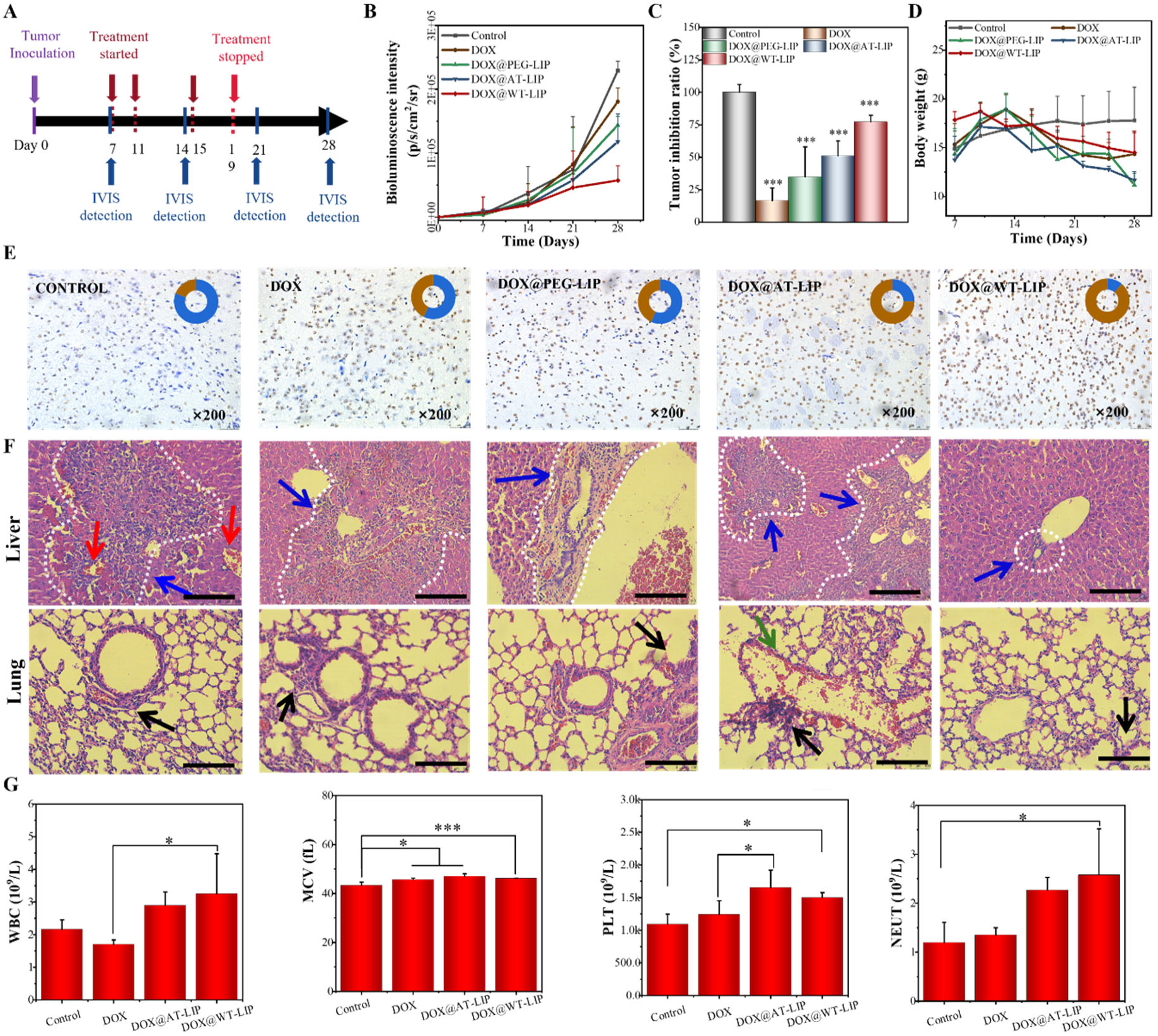
Antitumor efficacy on the U251 orthotopic GBM bearing mice model and H&E, blood biochemistry. (A) Schematic diagram of luciferase expressing orthotopic GBM model and drug treatment. After GCs implantation, tumor growth was monitored by IVIS imaging system; (B) Quantification of signals from the orthotopic GBM bearing mice; (C) The quantification signal of each group at the end of the experiment; (D) Body weight variation of U251-luc GBM-bearing mice during administration periods; (E) TUNEL staining U251-luc GBM-bearing brain collected at the day after last administration. (F) H&E staining of the U251 GBM-bearing livers and lungs collected at the day after last administration, the blue arrow represents necrosis site in liver and ductular proliferation, the black arrow represents cancer cells in lungs, the red arrow represents bleeding site; (G) Blood biochemistry of GBM model mice after administration with DOX@WT-LIP, DOX@AT-LIP, free DOX, and normal saline at an equal DOX concentration of 4 mg/kg. \**p* < 0.05. \*\*\**p* < 0.001

TUNEL staining was performed to assess apoptosis in tumor tissue (Figure 5E). Few TUNEL-positive (dark brown) apoptotic bodies were observed in the control group, whereas more apoptotic cells were observed in the free DOX, DOX@PEG-LIP and DOX@AT-LIP groups. The DOX@WT-LIP group showed a higher density of TUNEL-positive cells within the tumor region, consistent with increased apoptosis following treatment.

### 2.10 Systemic Toxicity

Major organs were collected for H&E staining, showing liver necrosis/ductular proliferation across groups but with less pronounced changes in the DOX@WT-LIP group; the DOX@AT-LIP group additionally exhibited liver injury and focal pulmonary hemorrhage (Figure 5E), while no clear abnormalities were noted in other organs. Routine hematology at 24 h (Figure 5G, Supplementary Figure 3B) indicated DOX-associated changes (decreased WBC and increased MCV), whereas DOX@WT-LIP largely preserved RBC-related indices with WBC within the normal range. In contrast, DOX@AT-LIP increased MCV and elevated PLT above the normal range, suggesting a less favorable hematologic profile.

## 3 Conclusion

In this study, we developed doxorubicin-loaded liposomes co-modified with the cell-penetrating peptide TAT and the dual-targeting peptide WTW (DOX@WT-LIP), guided by the CRITID cascade delivery concept for glioblastoma (GBM) nanomedicine. This innovative design led to significant improvements, including enhanced blood-brain barrier (BBB) transport (up to 60.74% *in vitro* transport efficiency), increased accumulation in GBM cells (DOX@WT-LIP uptake was 1.28-, 1.76-, and 2.12-fold higher than DOX@AT-LIP, DOX@PEG-LIP, and free DOX, respectively, in U251-MG cells), and superior apoptosis-related antitumor activity. These findings support a nuanced transport mechanism in brain endothelial cells (BCECs) and a targeted binding/uptake process in glioma cells (GCs), influenced by the synergistic action of WTW and TAT. The resulting higher drug exposure in the brain and an elevated GBM-to-brain (G/B) ratio correlated with improved tumor inhibition (74.92%) while potentially limiting exposure to normal brain tissue. Furthermore, reduced drug distribution in major organs contributes to lower systemic toxicity, collectively underscoring the promising potential of DOX@WT-LIP as an effective GBM drug delivery strategy.

## Acknowledgements

This work was supported by the National Key R&D Program of China (2022YFC2405703). We appreciate Dr. Bo Gao from the Analytical & Testing Center of Sichuan University for helping with MS characterization.

## Data and materials availability

All data are available in the main text or the supplementary materials.

