## Supplementary for "Cascaded Dual-Targeting (WTW/TAT) Liposomes Delivery DOX for Precision Glioblastoma Treatment"

### 1. Materials

#### 1.1 Materials and Reagents

WTWEYTK-Cys (WTW), Angiopep-2-Cys (Ang), and Cys-TAT (TAT) peptide were synthesized by Biotech BioScience & Technology (Shanghai, China). Doxorubicin (DOX) was purchased from Meilun Biotechnology (Dalian, China). Soybean phospholipid (SPC), cholesterol (CHO), DSPE-PEG$_{2000}$, and DSPE-PEG$_{2000}$-Mal were provided by Aladdin Bio-Chem Technology (Shanghai, China). Co6 was purchased from Sigma-Aldrich (Saint Louis, MS). DiD was purchased from Beyotime Biotechnology (Shanghai, China). Primary antibodies and secondary antibodies (anti-γ-H2AX, anti-CD31) were purchased from Signalway Antibody (Maryland, USA). MTT was purchased from J&K Scientific Ltd. (Beijing, China). Lyso Traker Red was purchased from UElandy. Inc. (Suzhou, China). ER Tracker Red was provided by MaoKang Biotechnology Co., Itd. (Shanghai, China). DAPI was bought from Solarbio Science & Technology (Beijing, China). D-luciferin potassium salt was obtained from BioVision, Inc. (U.S.A.). TUNEL Apoptosis Assay Kit was purchased from Roche (Mannheim, Germany). All other reagents were of analytical grade.

#### 1.2 Cells and Animals

The human glioblastoma (U251-MG) cells, mouse brain endothelial (bEnd.3) cells, and mouse astrocytes (M1800-57) cells were purchased from Jennio Biotech. Co., Ltd. (Guangzhou, China). RAW264.7 cells was purchased from Cyagen Biosciences Inc. (Suzhou, China). Mouse fibroblasts (L929) Cells, human glioblastoma (U87) cells, human epithelial ovarian cancer (SKOV-3) cells, and human umbilical vein endothelial (HUVEC) cells were provided from Institute of Biochemistry and Cell biology (Shanghai, China). Cells were maintained in DMEM medium which contained 10% (v/v) FBS, and 1% (v/v) penicillin/streptomycin solution. Cells were all cultured in a humidified atmosphere (Thermo Scientific, Waltham, MA, U.S.A.) of 5% $CO_2$ at 37°C. Male SPF Balb/c nude mice (6~8 weeks) were purchased from Chengdu Dossy experiment Animals Co., Ltd. All animal experimental protocols were approved and conducted by the Animal Ethics Committee of the Institutional Animal Care and Use Committee of Sichuan

University and performed following the Guidelines for Care and Use of Laboratory Animals of Sichuan University.

### 2. Experimental Section

#### 2.1 Synthesis and characterization of the conjugates

DSPE-PEG$_{2000}$-WTW was synthesized by conjugating the terminal maleimide (Mal) of DSPE-PEG$_{2000}$-Mal and the sulfhydryl group of cysteine of WTW as the previous method.[1] Briefly, WTW, DSPE-PEG$_{2000}$-Mal, and Triethylamine (molar ratio of 1:1:3) were dissolved in the mixture of trichloromethane and methanol (volume ratio of 2:1) with slow stirring in darkness for 24 hours under Nitrogen gas protection. The mixture was evaporated (SY-2000, Yarong, China), dialyzed (MEMBRS-CELL, MWCO = 3500 Da) in DI water and frozen dried to get DSPE-PEG$_{2000}$-WTW. Similarly, DSPE-PEG$_{2000}$-Ang and DSPE-PEG$_{2000}$-TAT were synthesized by the same method. The structure of DSPE-PEG$_{2000}$-WTW/Ang/TAT was verified by $^1$H NMR (600 MHZ, Unity Inova spectrometer, Phoenix, USA), and MALDI-TOF-MS (Thermo Fisher Scient, Germany).

#### 2.2 Preparation and characterization of liposomes

##### *2.2.1 Preparation of liposomes*

DOX-loaded liposomes were prepared using the classical method (Y) of transmembrane pH gradient.[2] To synthesize WTW and TAT co-modified DOX-loaded liposome (DOX@WT-LIP), SPC (0.59 mmol), CHO (0.36 mmol), DSPE-PEG$_{2000}$ (0.015 mmol), DSPE-PEG$_{2000}$-WTW (0.025 mmol), and DSPE-PEG$_{2000}$-TAT (0.01 mmol) were sufficiently dissolved in a 50 mL pear-shaped bottle containing trichloromethane/methanol solution (volume ratio of 2:1), and the mixture was then evaporated under vacuum (0.09 kPa) to form a lipid membrane. Next, the citric acid solution (4 mL, 0.3 mol/L, pH 4.0) was added into the lipid membrane to hydrate at 37°C for 40 min. The mixture was ultrasonic, rehydrated for 10 min and then was filtered through a polycarbonate filter with a pore size of 100 nm (Avanti Mini-Extruder Set, Avanti Polar Lipid Inc, USA) to obtain blank liposome. Then, the blank liposome was adjusted the pH of the external phase to 7.2 with $Na_2CO_3$ solution (0.5 mol/L, pH 11.2), and then incubated with DOX (the weight ratio of lipid and DOX was 20:1) in a shaker (100 rpm) at 50°C for 30 min. After cooling down to room temperature, free DOX was removed by centrifuging with a tubular ultrafiltration membrane (100,000 MW, Amicon Ultra4, Millipore Inc. U.S.A.). The derivates were all synthesized as the above procedure described, including PEG-modified DOX-loaded liposome (DOX@PEG-LIP), Angiopep-2 and TAT-modified DOX-loaded liposome (DOX@AT-LIP). Formulations of liposomes were shown in Table 1.

**Supplementary Table 1. Compositions of liposomes (mol)**

| Sample | SPC | CHO | Functional material compositions |
|---|---|---|---|
| PEG-LIP-DOX (DOX@PEG-LIP) | 59% | 36% | 5% DSPE-PEG$_{2000}$ |
| Ang-TAT-LIP-DOX (DOX@AT-LIP) | 59% | 36% | 1.5% DSPE-PEG$_{2000}$, 1% DSPE-PEG$_{2000}$-TAT, 2.5% DSPE-PEG$_{2000}$-Ang |

| | | | |
|---|---|---|---|
| WTW-TAT-LIP-DOX (DOX@WT-LIP) | 59% | 36% | 1.5% DSPE-PEG$_{2000}$, 1% DSPE-PEG$_{2000}$-TAT,<br>2.5% DSPE-PEG$_{2000}$-WTW |

***2.2.2 Co6-loaded liposomes***

Briefly, to synthesize WTW and TAT-modified Co6-loaded liposomes, SPC, CHO, DSPE-PEG$_{2000}$, DSPE-PEG$_{2000}$-TAT, and DSPE-PEG$_{2000}$-WTW (molar ratio of them was shown in Table S1) were all dissolved in a trichloromethane/methanol (volume ratio of 2:1) solution, and Co6 (Co6 and lipid were in a weight ratio of 1:20) was added into the mixture. The mixture was then evaporated under vacuum to form a thin dry lipid membrane. Next, the lipid membrane was hydrated with PBS buffer at 37°C for 40 min, extruded through a polycarbonate filer of 100 nm pore size, and then centrifuged in a tubular ultrafiltration membrane at 8500 rpm for 25 min. Similarly, PEG-modified Co6-loaded liposome (Co6-PL), Angiopep-2 and TAT-modified Co6-loaded liposome (Co6-AT-LIP) were synthesized as the procedure mentioned above, respectively.

***2.2.3 Characterization of liposomes***

The hydrodynamic diameter size, ζ-potentials, and PDI of the liposomes were measured using a dynamic light scattering analyzer (Nano ZS90, Malvern Instruments Ltd., U.K.) at 25°C. Measurements were made in triplicate. The nuclear-shell structures of liposomes were observed by a transmission electron microscope (TEM) and a cryo-transmission electron microscope (cryo-EM). To image the liposome by TEM, 10 μL liposome suspension was first deposited on a glow-discharged carbon-coated grid. After drying at room temperature, a drop of 3% phosphotungstic acid was added to the grid. The liposome was then dried and observed under TEM (JEM-100CX, JEOL, Tokyo, Japan). For cryo-EM imaging, 3 μL liposome suspension was added on glow-discharged (45 s) 200-mesh R2/1 Quantifoil Cu grids. The grids were blotted for 3 s in 100% humidity without blot offset and rapidly froze in liquid ethane using Vitrobot Mark IV (Thermo Fisher). The frozen grids were loaded into a Talos Arctica (Thermo Fisher) with an operating voltage of 200 kV, a condenser lens aperture of 50 μm, and a spot size of 7. The liposomes were observed at a microscope magnification of 150,000×.

**2.3 In vitro drug loaded and drug release behavior of liposomes determination**

The drug encapsulation efficiency (EE) and loading efficiency (LE) of liposomes were calculated according to the following formula (1) and (2), respectively. $M_{\text{DOX in liposomes}}$ represented the mass of DOX in liposomes, $M_{\text{total DOX}}$ represented the total mass of DOX, $M_{\text{lipid}}$ represented the mass of lipid.

$$EE\% = M_{\text{DOX in liposomes}} / M_{\text{total DOX}} \times 100\% \quad (1)$$

$$LE\% = M_{\text{DOX in liposomes}} / (M_{\text{total DOX}} + M_{\text{lipid}}) \times 100\% \quad (2)$$

For drug release in vitro, DOX loading liposomes and free DOX (500 μL, DOX-equivalent 5 mg/mL) were added into dialysis bags (3,500 MW) and then against 20 mL PBS buffer with shaking (40 rpm) at 37°C for 48 h. The release medium (2 mL) was sampled at the set time points, replaced with an equal volume of PBS buffer, and measured by a UV–vis spectrometer (U-3010, Hitachi, Japan) at 450 nm. The formula for calculating the drug release ratio (DRR) was shown in (3), the $\text{Dose}_{\text{released DOX}}$ represented the amount of

DOX dose in the release medium, the $Dose_{total\ DOX}$ represented the total DOX dose in the liposome.

$$DRR\ (\%) = Dose_{released\ DOX} / Dose_{total\ DOX} \times 100\% \qquad (3)$$

**2.4 Hemolysis test**

The hemolysis ratios of DL, TDL, ADL, WDL, ATDL, and WTDL were evaluated. PBS buffer and 1% (v/v) Triton solution were set as negative control and positive control, respectively. Firstly, the whole blood of the rabbit was diluted with PBS buffer ($V_{Blood} : V_{PBS} = 3: 6$), centrifuged at 1500 rpm for 8 min, decanted the supernatant, and added fresh PBS buffer. The process was repeated five times to obtain a 10% (v/v) red blood cell (RBC) suspension. Subsequently, 500 μL DOX-loaded liposomes (DOX-equivalent 20 μg/mL) were mixed with an equal volume of 10% RBC suspension and incubated with shaking (180 rpm) at 37°C for 2 h. The mixture was then centrifuged at 1500 rpm for 8 min, and the supernatant was measured by microplate spectrophotometer 3350 (Tunable VERSA max, Molecular Devices, USA) at 545 nm. The hemolytic ratio (HR%) was calculated by the formula (4), $OD_{sample}$, $OD_{negative}$, and $OD_{positive}$ represented the absorbance of supernatant in different groups, respectively.

$$HR\ (\%) = (OD_{sample} - OD_{negative}) / (OD_{positive} - OD_{negative}) \times 100\% \qquad (4)$$

**2.5 Cytotoxicity of liposomes against different types of cells**

MTT was used to determine the cytotoxicity of liposomes to cells. U251-MG, bEnd.3, and L929 cells were seeded into 96-well plates at a density of $1\times10^4$ cells/well for 24 h, respectively. Cells were then incubated with WT-LIP, and its derives (1 μg/mL) for 48 h. The control group was incubated with a DMEM medium. Next, the medium was replaced with 200 μL fresh mixture (DMEM containing 10% MTT) for incubation 4 h. The supernatant was discarded and washed by PBS. 150 μL DMSO was then added into the well to dissolve the purple formazan crystals by shaking 15 mins. The absorbance intensity was measured by a plate reader at 450 nm. The cell viability (%) was calculated using the formula (6), where $OD_{test}$ and $OD_{control}$ represented the absorbance of cells treated with different liposomes and DMEM medium, respectively.

$$Cell\ viability\ (\%) = OD_{test} / OD_{control} \times 100\% \qquad (5)$$

Subsequently, the inhibition effect of DOX@WT-LIP and its derives *in vitro* was investigated. U251-MG cells were seeded into 96-well plates at a density of $1\times10^4$ cells/well for 24 h. DOX@WT-LIP, its derives and free DOX were added into wells at the final testing concentrations (equivalent to DOX: 0~20 μg/mL) to incubate different times (24, 48 h for U251-MG cells). The control group was incubated with a DMEM medium. The medium was then replaced with a mixture of MTT solution (20 μL, 5 mg/mL) and 180 μL fresh medium to incubate at 37°C for 4 h. Afterwards, the mixture was replaced with 150 μL DMSO to dissolve the purple formazan crystals. The absorbance intensity in each well was measured by microplate spectrophotometer 3350 at 480 nm. The cell viability (%) was calculated using the formula (6).

**2.6 Cellular Uptake**

Firstly, U251-MG, bEnd.3, M1800-57, SKOV3, L929, RAW 264.7, and HUVEC cells were seeded into glass bottom plates at a density of $1\times10^5$ cells/plate for 24 h, respectively. Afterwards, cells were co-incubated with Co6-PL, Co6-ATL and Co6-WTL (10 μg/mL) for 4 h, washed with PBS twice, and then fixed with 4% paraformaldehyde. The nuclei were stained with DAPI for 10 min. Finally, the cells were washed with PBS again and observed under confocal microscopy (Zeiss LSM 880, Germany) using 405, and 488 nm wavelength channels.

Cellular uptake was quantitatively analyzed with average fluorescence intensity by flow cytometry. U251-MG, bEnd.3, M1800-57, SKOV3, L929, RAW 264.7, and HUVEC cells were seeded into 12-well plates at a density of $1\times10^5$ cells/plate for 24 h, respectively. The cells were incubated with Co6-PL, Co6-ATL, and Co6-WTL in a concentration of 10 μg/mL for 4 h, washed with PBS twice, digested, centrifuged at 800 rpm for 3 min, and resuspended with 500 μL fresh PBS. Then the cells were transferred to a flow cytometry tube and measured by flow cytometry (Moflo XDP, Beckman Coulter, Brea, U.S.A.).

**2.7 Endocytosis behavior and lysosome/ER colocalization exploration**

***2.7.1 Endocytosis behavior Exploration***

U251-MG, and bEnd.3 cell were seeded into a 12-well plated at a density of $1\times10^5$ cells/plate for 24 h, respectively. The medium was replaced with fresh medium and subsequently five endocytosis inhibitors (chlorpromazine, genistein, amiloride, methyl-β-cyclodextrin, or cytochalasin D) were separately added to the medium at concentrations of 20, 100, 20, 20, or 5 μg/mL for 1 h, respectively. Similarly, the low temperature groups were incubation under 4℃ for 1 h. After preincubation with inhibitors, the cells were further incubated with Co6-PL, Co6-ATL, and Co6-WTL (500 μL, 100 μg/mL) for 3 h. Cells were then washed, digested, and collected for analysis by flow cytometry (Moflo XDP, Beckman Coulter, Brea, U.S.A.).

***2.7.2 Lysosome/ER colocalization***

U251-MG, and bEnd.3 cell were seeded into glass bottom plates at a density of $1\times10^5$ cells/plate for 24 h, respectively. The medium was replaced with 500 μL Co6-WTL for 4 h incubation. Cells were washed with PBS, co-incubated with Lyso Tracker Red or ER Tracker Red for 30 min, and the nuclei were stained with DAPI for 20 min. Cells were then washed, added a drop of glycerin to keep color, and observed using CLSM (Zeiss LSM 880, Germany).

**2.8 Expression of γ-H2AX in tumor cells measurement**

To investigate the antitumor mechanism of liposomes in GCs, U251-MG cells were incubated with DOX-loaded liposomes ad then observed the expression of γ-H2AX and the changes of cell morphology. U251-MG cells were incubated with 500 μL DOX@PEG-LIP, DOX@AT-LIP, DOX@WT-LIP, and free DOX (DOX-equivalent 20 μg/mL) for 0.5 or 24 h. Cells were then washed with PBS twice, fixed with 4% paraformaldehyde for 15 min, incubated with 0.1% Triton X-100 for 10 min. Next, cells were blocked by 3% (w/v) bovine serum albumin (BSA, Aladdin, China) for 1 h, and incubated with primary antibody **γ-**H2AX overnight at 4°C. Cells were washed with PBS twice, incubated with secondary antibody Alex-488 for 1 h, and then stained by phalloidin for 1 h. The nuclei were stained with DAPI. Cells were observed by CLSM (Zeiss LSM 880, Germany) fluorescent filter sets: Alex-488 (Ex: 495 nm, Em: 519 nm), phalloidin (Ex: 540 nm, Em: 565 nm), DAPI (Ex: 340 nm, Em: 488 nm).

**2.9 Evaluation transport efficiency of liposomes *in vitro* BBB/GBM model**

#### *2.9.1 Establishment of BBB/GBM model in vitro*

In vitro BBB model was established according to the previous report.[3] Briefly, bEnd.3 cells were seeded in the upper chamber of the 12-well transwell (Corning, NY, USA) at a density of $1\times10^5$ cells/well and consistently cultured for 14 days to form the bEnd.3 monolayers. BBB models *in vitro* were monitored for the changes of the transendothelial electric resistance (TEER) by Millicell ERS (Millipore, USA) in 14 days.

To establish *in vitro* GBM model, bEnd.3 cells were seeded into upper chamber of 12-well transwell for 14 days. U251-MG cells were seeded in another 12-well at a density of $1\times10^5$ cells/well overnight. The upper chambers containing the bEnd.3 monolayers were then transferred into the 12-well plates containing U251-MG cells to form the *in vitro* GBM model.

#### *2.9.2 Evaluation on transport efficiency of liposomes via in vitro BBB model*

The bEnd.3 monolayers were constructed in upper chamber of 12-well transwell cell culture plate according to the procedures described. 500 μL DMEM containing DL, ATDL, WTDL, and free DOX (DOX-equivalent of 40 μg/mL) were added into the upper chamber, and then 1mL PBS buffer was introduced into the lower chamber of the 12-well transwell plate. The PBS buffer in the lower chamber was collected 100 μL every 10 min, and then replaced with another equal volume of fresh PBS buffer until 480 min. The concentration of DOX was determined by high-performance liquid chromatography (HPLC). The permeated DOX dose arrived in lower chamber (PD%) was calculated by the formula (7), $DOX_{test}$ and $DOX_{total}$ represented the amount of DOX reaching the lower chamber and total DOX dose, respectively.

$$PD\ (\%) = DOX_{test} / DOX_{total} \times 100\% \qquad (6)$$

#### *2.9.3 Confocal microscopy observation on transcytosis via GBM model in vitro*

The *in vitro* GBM model was constructed according to the procedures described above. Briefly, the bEnd.3 monolayers were constructed in 12-well transwell cell culture plates for incubation 14 days. U251-MG cells were seeded in another 12-well plate at a density of $10^5$ cells/well for 24 h. The upper chamber with bEnd.3 monolayers were then transferred into the 12-well plates containing U251-MG cells.

Subsequently, DL, ATDL, WTDL, and free DOX (500 μL, DOX-equivalent 40 μg/mL) were added into the upper chamber to co-incubate for 24 h, and then 1 mL DMEM was added into the lower chamber. Afterwards, the bEnd.3 monolayers in the upper chamber were removed, washed with PBS twice, fixed with 4% paraformaldehyde for 15 min, added 0.1% Triton-100 for 20 min, and then blocked by 3% BSA solution for 1 h. Next, the monolayers were stained by phalloidin for 1 h (1:200 dilution), and the nuclei were stained by DAPI for 10 min. After washing with PBS twice, the bEnd.3 monolayers were then transferred on glass microscope slides with a drop of glycerin to keep them fresh, and observed the 3D images by CLSM (Zeiss LSM 880, Germany).

#### *2.9.4 Flow cytometry analysis on transcytosis via GBM model in vitro*

To establish *in vitro* GBM model, the bEnd.3 monolayers were incubated in a 12-well transwell cell culture plate for 14 days, and U251-MG cells were cultured in another 12-well plate for 24 h before the monolayer filmed formation. Subsequently, DL, ATDL, WTDL, and free DOX (500 μL, DOX-equivalent 40 μg/mL) were introduced into the upper chamber, and 1 mL DMEM medium was added into the lower chamber. After co-incubation for 2 and 24 h, U251-MG cells in the lower chamber were washed with PBS twice, collected, resuspended with 500 μL PBS, and measured by flow cytometry (Moflo XDP, Beckman Coulter, Brea, U.S.A.).

***2.9.5 Live/Dead assay***

Cell Live/Dead staining was performed to compare the inhibitory effects of DOX-loaded liposomes against U251-MG cells. As the above mentioned, the bEnd.3 monolayers after incubation 14 days were transferred in another 12-well plate containing U251-MG cells to construct the GBM *in vitro* models. DL, ATDL, WTDL, and free DOX (500 μL, DOX-equivalent 40 μg/mL) were then added into the upper chamber for 24 h. U251-MG cells were then dyed with working solution A and B from the dye kit (LIVE/DEAD Cell Viability Kit, Thermo Fisher, USA). After incubating for 30 min, the cells were photographed with an inverted fluorescence microscope. The viable cells appeared green while the dead appeared red under blue excitation light.

**2.10 Evaluation of liposomes effect of targeting and inhibited *in vivo* GBM model**

***2.10.1 Establishment of orthotopic GBM xenograft***

The orthotopic GBM xenograft mice models were established by stereotaxic inoculation of luciferase-transfected U251-luc human glioblastoma cells.[4] Luciferin labeled U251-luc cells (U251-luc) are inserted the luciferase gene into cell chromosomes, which can be imaged the green fluorescence when oxidizing with D-luciferin potassium salt[5]. The size of GBM tumor was observed by injecting D-luciferin potassium salt into the GBM model mice and conducting tumor fluorescence imaging within 15 minutes by IVIS Spectrum imaging system.

Briefly, nude mice were anesthetized by an intraperitoneal injection with 1% sodium pentobarbital, intracranially injected U251-luc cells ($1\times10^6$ suspended in 5 μL DMEM) into the right striatum (1.5 mm lateral, 1.5 mm longitudinal, and 3 mm depth) using a specifically 24# trocar made propeller, and a brain stereotactic fixation device. Mice were raised under SPF standard conditions for one week and used for subsequent experiments. Following animal ethics, the mice were euthanized when the maximum weight loss approached 20%.

***2.10.2 In vivo biodistribution***

DiD-PEG-LIP (DiD-PL), DiD-Ang-TAT-LIP (DiD-ATL), and DiD-WTW-TAT-LIP (DiD-WTL) were prepared as the same method of Co6-WTL synthesized described above, wherein a DiD dose of 0.03 mg/kg for mice body weight. After one week raised, intracranial U251-luc GBM mice were randomly divided into three groups, and injected with DiD-PL, DiD-ATL, and DiD-WTL (100 μL, 10 μg/mL) through the tail vein. The fluorescence distribution of the whole body was then observed (excitation: 640 nm, emission: 670 nm) with the IVIS Spectrum imaging system (PerkinElmer, USA) at 2, 6, 12, and 24 h.

Subsequently, mice were euthanized with 20% urethane after injected liposomes 24 h, and major organs (heart, liver, kidney, spleen, lungs, and brain) were washed with 0.9% saline, and then photographed the fluorescence distribution using the IVIS Spectrum imaging system. The fluorescent quantitation of organs and brain tumors was calculated by using the IVIS Spectrum Software in the radiant efficiency model.

***2.10.3 Immunofluorescence observation of brain in GBM model***

The brain tissues were fixed in 4% paraformaldehyde for 24 h, dehydrated by 15% and 30% sucrose solution for 24 h in sequence, and the final paraffin-embedded and prepared into 10 μm ultra-thin sections. Then the sections of the brain were incubated with the primary anti-CD31 antibody at 4°C overnight, and the slices were stained with Alex 488-conjugated secondary antibody at room temperature for 2 h. Subsequently, the nuclei were stained with DAPI for 10 min. The slices were then added anti-fade mounting medium and observed the fluorescence images by CLSM (Zeiss LSM 880, Germany).

***2.10.4 Pharmacokinetics study***

SD mice were fasted for 6 h to empty the digestive tract. WTDL was administered through tail vein injection (DOX-equivalent, 4 mg/mL). The blood samples were collected by the orbital venous plexus of mice at different point (0, 1, 3, 6, 12, 24, and 48 h), and then transferred to EDTA-treated tubes and centrifuged at 12000 rpm for 15 min at 4℃. The blood serum was diluted with methyl alcohol (500 μL) and then thoroughly vortexed, ultrasonicated, and centrifuged at 5000 rpm for 5 min. 200 μL of supernatant was removed and concentrated for LC/MS analysis.

***2.10.5 Evaluation of antitumor in vivo***

U251-luc GBM-bearing Balb/c mice were established as previously mentioned and raised for one week. Mice were randomly divided into five groups. On the 7th, 11th, 15th, and 19th day after implantation, mice were injected with 50 μL WTDL, ATDL, DL, free DOX, and saline, and the administration dose of DOX was 4 mg/kg. Meanwhile, the body weights and overall survival were monitored for all groups.

The tumor progress was observed and semi-quantitatively analyzed by IVIS Spectrum imaging system within 15 min after intraperitoneal injection of D-luciferin potassium salt (100 μL, 150 mg/kg) on the 7th, 14th, 21st, and 28th day. Additionally, the mice were euthanized after fluorescence imaging on the 28th day, and brains and major organs were sampled for H&E staining, and brains were stained for TUNEL observation.

**2.11 Blood biochemistry analysis**

The U251-luc GBM-bearing mice were randomly divided into four groups (3 mice per group). At day 10/13/16/19, mice in each group were injected with WTDL, ATDL, free DOX, and normal saline, and the administration dose of DOX was 4 mg/kg. Whole blood samples were collected 24 h later after the last administration from the orbit. All blood samples were examined to obtain blood routine data.

**2.12 Statistical analysis**

Comparisons of data were performed by an independent sample t-test (two groups), and a one-way ANOVA (three or more groups). $p < 0.05$, 0.01, and 0.001 were considered a statistically significant difference and denoted with *p<0.5, ** p<0.1, and *** p<0.01, respectively.

**3. Results**

**3.1**

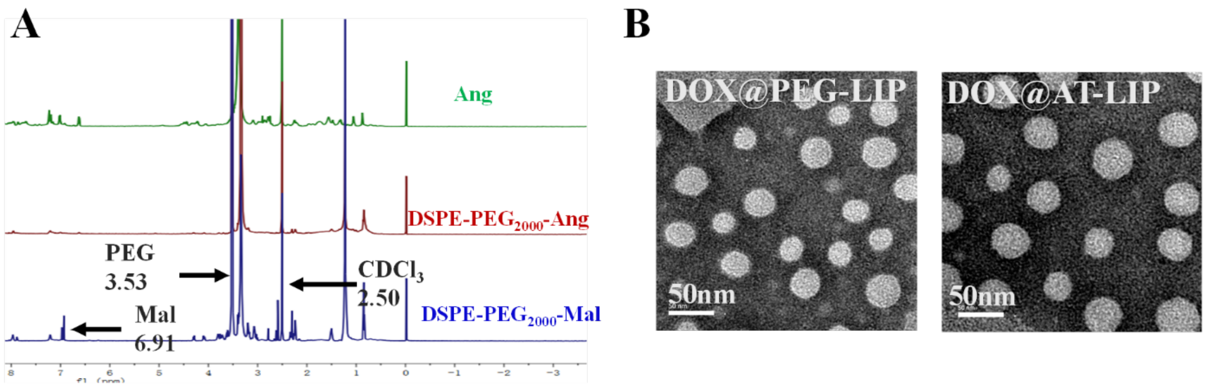


Supplementary Figure 1. (A) 1H NMR of DSPE-PEG2000-Ang; (B) cryo-EM image of DOX@PEG-LIP and DOX@AT-LIP.

**3.2**

**Supplementary Table 2. Physiochemical characteristics of DOX-loaded liposomes**

| Liposomes | PDI | LE (%) | EE (%) |
|---|---|---|---|
| DOX@PEG-LIP | 0.16 ± 0.01 | 2.96 ± 0.01 | 98.36 ± 0.48 |
| DOX@T-LIP | 0.16 ± 0.01 | 2.87 ± 0.03 | 98.38 ± 1.05 |
| DOX@A-LIP | 0.18 ± 0.01 | 2.44 ± 0.02 | 96.09 ± 0.83 |
| DOX@W-LIP | 0.20 ± 0.01 | 2.47 ± 0.30 | 91.55 ± 1.10 |
| DOX@AT-LIP | 0.10 ± 0.01 | 2.39 ± 0.03 | 96.92 ± 1.31 |
| DOX@WT-LIP | 0.26 ± 0.01 | 2.84 ± 0.08 | 94.17 ± 1.30 |

**3.3**

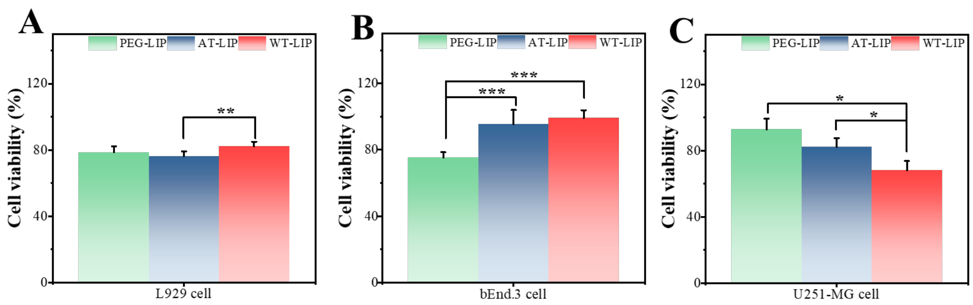


Supplementary Figure 2. The MTT assay of PEG-LIP, AT-LIP, and WT-LIP against L929 (A), bEnd.3 (B), and U251-MG cells (C) (*$p < 0.05$, **$p < 0.01$, ***$p < 0.001$; represented the significantly different with control group)

#### 3.4 Pharmacokinetic study

Supplementary Table 3 DOX pharmacokinetic parameters of DOX@WT-LIP

| Formulation | Dose (mg/kg, n=3) | $C_{max}$ (mg/L) | $T_{max}$ (h) | $T_{1/2}$ (h) | $AUC_{0-t}$ (mg h $L^{-1}$) | $MRT_{0-t}$ (h) |
|---|---|---|---|---|---|---|
| DOX@WT-LIP | 4 | 14.148 | 1 | 1.965 | 52.147 | 14.629 |

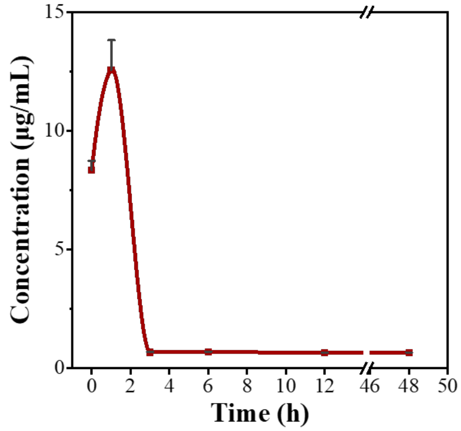


Supplementary Figure 3. DOX pharmacokinetic curve of GBM-targeted doxorubicin liposome DOX@WT-LIP

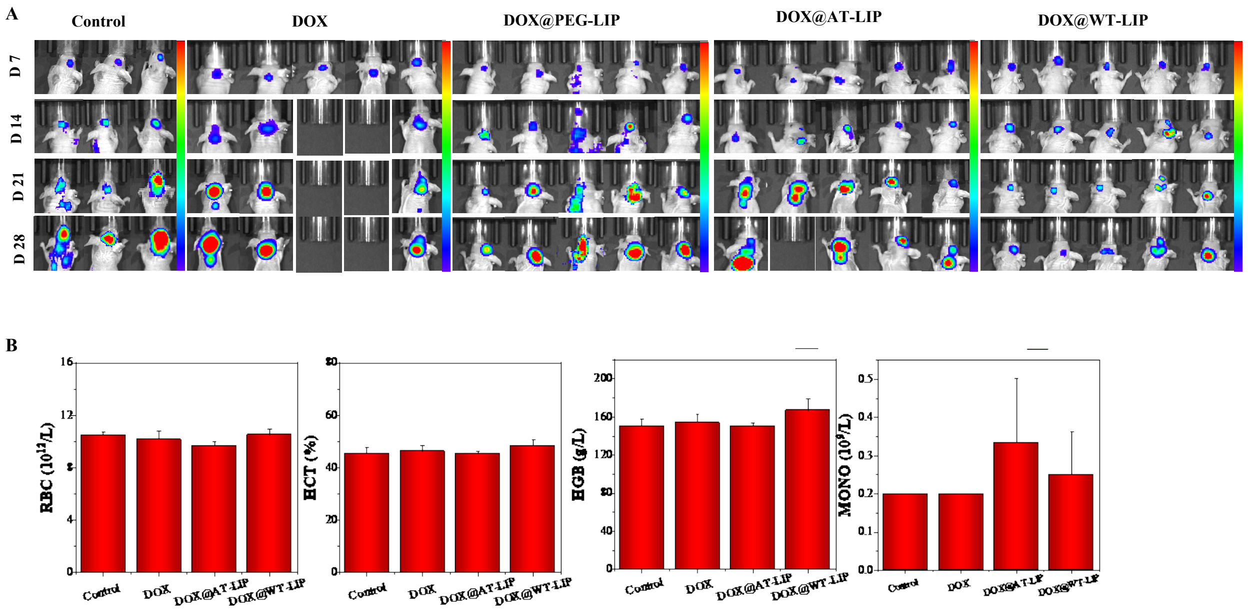


Supplementary Figure 4. (A) DOX loading liposomes efficacy in GBM mouse model. (B) Blood test after treatment.

Supplementary
